# Connectome Constrained Graphical Models of MEG Coherence

**DOI:** 10.1101/785600

**Authors:** Anirudh Wodeyar, Ramesh Srinivasan

## Abstract

Structural connectivity by axonal fiber bundles provides the backbone for communication between neural populations. Since axonal transmission occurs on a millisecond time scale, measures of M/EEG functional connectivity sensitive to phase synchronization in a frequency band, such as coherence, are expected to reflect structural connectivity. We develop a complex-valued Gaussian Graphical Model (cGGM) of MEG coherence whose edges are constrained by the structural connectome. The cGGMs’ edge strengths are summarized by partial coherence, a measure of conditional dependence. We made use of the adaptive graphical lasso (AGL) to fit the cGGMs which allows us to perform inference on the hypothesis that the structural connectome is reflected in MEG coherence in a frequency band. In simulations, we demonstrate that the structural connectivity’s influence on the cGGM can be inferred using the AGL. Further, we show that fitting the cGGM is superior to alternative methods at recovering the structural connectome. Graphical modeling of MEG coherence is robust to the source localization estimates required to map MEG from sensors to the cortex. Finally, we show how cGGMs can be used to explore how distinct parts of the structural connectome contribute to MEG coherence in different frequency bands. We think the cGGM is a useful tool that can improve interpretation of MEG coherence by making a direct link to the structural connectome.

## 1. Introduction

Electrophysiological signals are sampled on a millisecond time scale capturing aggregate synaptic activity from populations of neurons. These neurophysiological signals have intrinsic time scales of dynamics, usually organized in terms of frequency bands that are distinctly modulated in different cognitive tasks and clinical disease states [1, 2, 3, 4, 5, 6]. Functional connectivity (FC) refers to statistical dependence between signals recorded from two different areas of the brain usually measured in a pre-defined frequency band. This broad definition encompasses different pre-processing methods and statistical models that emphasize different temporal and spatial scales of the underlying brain activity.

A widely used measure of functional connectivity expecting linear relationships and measuring millisecond time scale synchronization is coherence [7]. Coherence measures the consistency of relative phase and relative amplitude within a frequency band between signals recorded from two distinct areas. Coherence has long been interpreted as reflecting signal transmission with consistent axonal delays along white-matter tracts (structural connections) [8, 9, 10]. By extension, we would expect that between pairs of regions without structural connections, there would be no consistent relative phase and amplitude, and thus, low coherence. However, if a multi-step path over several structural connections persists between a pair of brain regions not connected by a direct structural connection, we may still expect non-zero coherence between them [11, 12, 13].

Coherence appears to capture something important about the communication patterns of the brain [14]. But, coherence makes it difficult to intepret the underlying communication pattern - which areas communicate with one another via a direct connection and which communicate indirectly via a multi-step path across intermediate nodes? Coherence is computed from the cross-spectral density which is the covariance structure for complex-valued data. If we assume, as is common, that the cross-spectral density fully parameterizes the structure of the data (i.e. we assume there are no nonlinear relationships and the data is narrow-band), then we can use a simple transformation of the cross-spectral density, by taking its inverse, that makes it easier to infer which connections are direct [15, 16]. The inverse of the covariance is a matrix termed the precision. This matrix represents the strength of linear relationships between a pair of brain areas when accounting for their relationships with all other brain areas [17, 18, 19]. As precision removes indirect covariance, we expect that the structural connectome maps better to the precision rather than to the coherence.

In electrophysiological data, we can estimate the coherence fairly easily. While it may be noisy when we have very few samples, estimation is straight-forward because the coherence is simply the normalized frequency-domain covariance (cross-spectral density). In contrast, taking the inverse of a matrix, as is necessary to estimate the precision and its equivalent normalization - partial coherence, is made difficult when we have a rank-deficient matrix as is usually observed with M/EEG (magneto-/electroencephalography). As we hypothesize that the precision is potentially constrained by the structural connectome and that to estimate the inverse we need regularization, we use a graphical lasso technique (which uses an L1 penalization). A critical innovation in this study is to modify the graphical lasso to allow us to use the *structural connectome* to guide the L1 penalization. This approach solves many problems with current techniques and provides an easier to interpret connectivity estimate that can help spur hypothesis-driven research.

In this study we develop a model of MEG functional connectivity from the structural connectome in the form of a complex-valued Gaussian Graphical Model (cGGM) and then show how we can estimate the parameters of this model using an adaptive graphical lasso (AGL). We use simulations to test this approach and to compare it to several commonly used methods of estimating functional connectivity demonstrating the advantages of our method irrespective of sample size. AGL estimation of a cGGM also performs better when the signals generated from the structural connectome are blurred due to forward simulation to sensors and source localization. Finally, we apply this method to resting-state MEG data where we show that functional connectivity in different frequency bands can be modeled by different connectivity patterns over the structural connectome.

## 2. Methods

### 2.1. Overview

This work is guided by the intuition that the statistics of neural activity data collected at the meso (ECoG) and macro (M/EEG) scales are constrained by the structural connectivity. As such, we have built a minimal generative computational model that is derived from the structural connectivity and developed a method by which we are able to perform inference on this model. We developed a complex-Gaussian graphical model (cGGM) of a widely used summary statistic used in EEG and MEG studies [7, 9, 20] - the coherence. We allowed the structural connectivity to guide the estimation of the cGGM and developed new simulations to link this work with M/EEG and ECoG data.

### 2.2. Modeling Framework

#### 2.2.1. Structural Connectome

We developed a model of the structural connection (SC) from a probabilistic atlas. We used streamlines generated with deterministic tractography by [21] using the HCP842 dataset [23] transformed to the MNI152 template brain obtained from FSL. In this dataset experts vet the streamlines to remove potentially noisy estimates of axonal fibers. We applied the Lausanne parcellation [22] of 114 cortical and 15 subcortical ROIs to the MNI152 template brain and generated a volumetric representation for each region of interest using the *easy lausanne* toolbox [24]. Each streamline was approximated by a single 100-point cubic spline using code adapted from the along-tract-stats toolbox [25]. By identifying the streamlines which terminated in a pair of ROIs (regions of interest) we were able to create the structural connectome (SC) for the Lausanne parcellation. Each streamline only connected a single pair of ROIs. An edge *W_ij_* for ROIs *i* and *j* existed if there was a streamline connecting the pair.

From this process, we built the 129 ∗ 129 undirected and unweighted structural connectome with 1132 edges. We reduced this matrix to 114 ∗ 114 with 720 edges (see Figure 1) after removing all the subcortical structures and limiting inter-hemispheric connections to homologous white-matter tracts. This latter step helped remove potentially noisy estimates of connections (while potentially increasing false negatives) where streamlines intersected and passed outside the cortical surface before reaching the terminal point in a brain region. The resulting template of structural connectivity shown in Figure 1 is referred to as the structural connectome (SC). This template is incomplete in that it does not include subcortical to cortical projections. Thus, functional connectivity resulting of structural connections not captured by this template may exist in the data. Our estimation procedure for the graphical models of functional connectivity described in the next section allows for such connections, if needed, to account for the statistical structure in the data.

**Figure 1:**
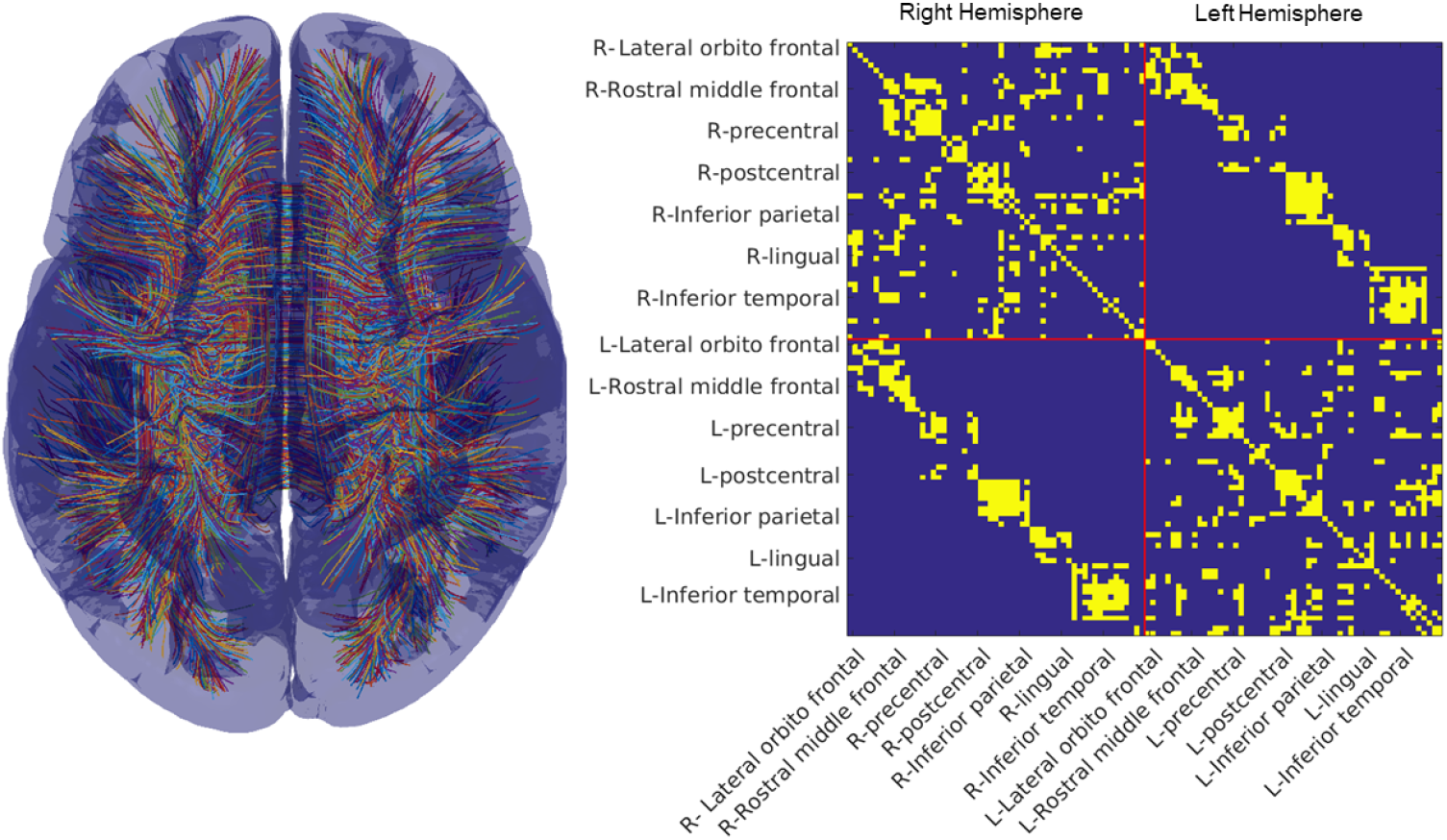
Structural Connectome. (Left) We show streamlines derived from the work of [21] using the HCP-842 dataset. (Right) The structural connectome for the 114 areas of the Lausanne parcellation. We have labeled a subset of areas each with 1 to 3 subdivisions (see [22] for all subdivisions of the Lausanne parcellation). We show the undirected and unweighted SC, with any non-zero edge being shown in yellow.

#### 2.2.2. Complex-Valued Gaussian Graphical Model (cGGM)

We assume that a vector of activity (**Z**) in one frequency band is a sample drawn from a **complex-valued** multivariate Gaussian (where Φ is the precision and is determined by the SC):

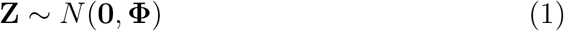

In the frequency domain, a signal can be characterized by samples of amplitude and phase, or equivalently, by complex valued coefficients with real and imaginary parts corresponding to sine and cosine components of the signal.

The complex-valued multivariate Normal for a zero-mean (where *E*(**Z**) = 0 + 0*i*) complex-valued Gaussian process [26] is defined as:

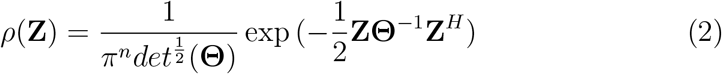

where

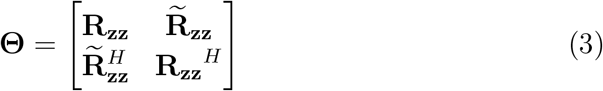

and

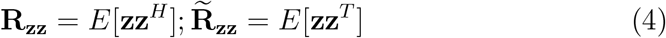

The key parameter in this model is the covariance matrix **Θ** and its inverse, the *precision* matrix **Φ** = **Θ**^*−*1^. As defined in equations 3 and 4, the covariance matrix for complex-valued data is composed of the familiar cross-spectrum **R_zz_** and the complementary cross-spectrum 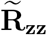. Most spectral analysis methods only make use of **R_zz_** and implicitly assume circular symmetry, i.e., 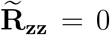 [26]. In this case, the complex-valued data is labeled as *proper*. With the assumption of circular symmetry, we can parameterize the complex-valued Gaussian using the precision defined as the inverse of the cross-spectrum, **Φ** = **Θ**^*−*1^ = **R_zz_**^*−*1^.

Each value in the precision matrix **Φ** is the conditional covariance between any two variables (here, sources representing two ROIs) given the other variables (all other ROIs). The precision represents a model of functional connectivity - the *conditional dependence* between sources. The strength of the conditional dependence represents the linear relationship between any pair of sources when linear effects from all other sources are removed (see Section 2.2.2 of [27] for an intuitive explanation in terms of multivariate linear regression). For any pair of sources, if the precision is zero, there is no need for a relationship between the sources to account for observed coherence. Such apparent coherences arise from connections mediated via other sources in the model. Note that the precision directly represents the complex-valued Gaussian Graphical Model or cGGM, a terminology we borrow from Whittaker [17]. We use the term “precision” and “cGGM” equivalently in the rest of the manuscript.

In the generative model, the precision matrix **Φ** has a nonzero entry only at edges that have a connection in the SC. We are assuming that in each frequency band, *coherence represents the result of joint random fluctuations of a set of oscillators whose connections are determined by the SC*. When performing inference on this model to estimate the precision vales, we use the graphical lasso in a cross-validated procedure allowing it to use the SC as a guide for the L1 penalization, if it reduces cross-validated error. In this way the non-zero locations and values of the precision are determined by the data.

### 2.3. Model Estimation

#### 2.3.1. Adaptive Graphical Lasso

The graphical lasso [28] is a method that has been applied in multiple fields in the past decade, from genomics [29], to fMRI functional connectivity [30, 31, 32] and climate models [33]. It is used to identify a sparse approximation to the precision matrix while solving problems arising from rank-deficiency and small numbers of samples. To apply the lasso, we optimize the penalized likelihood function for a multivariate Gaussian [34] to estimate the precision (where Θ is the cross-spectral density):

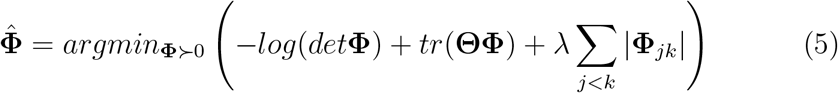

The penalization parameter *λ* in the graphical lasso determines the non-zero set of precision values. The output of the lasso from Equation 5 is the precision matrix 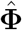, which represents directly the complex-valued Gaussian Graphical (cGGM).

We made use of the lasso to estimate the precision while taking advantage of the knowledge of the SC to hypothesize the likely locations of nonzero precision values. We made use of the lasso optimization from QUIC [35] using a matrix penalty term (this process is also called the adaptive lasso - [36]) determined by the SC with edges *W* (and *λ*_1_ = *λ*_2_):

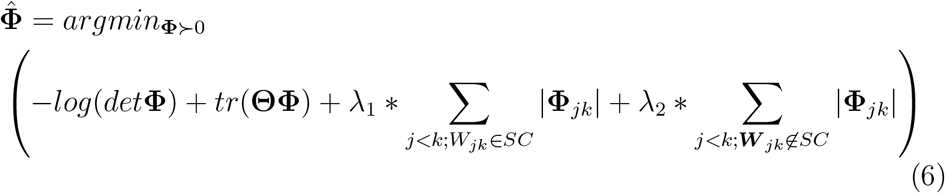

Note that in the limiting case of *λ*_1_ = *λ*_2_, the likelihood function is the same as it is for the graphical lasso. We determine the *λ*_1_ and *λ*_2_ using cross-validation. This crucial setup simultaneously provides a potential validation of the usefulness of the SC as a hypothesis on MEG functional connectivity while also serving as a principled thresholding mechanism for weak connections.

Past work applying a matrix penalty term for the graphical lasso [37] have used the SC connection weights directly on real-valued MEG signals to determine the penalization weighting, which imposes more structure on the penalization than we assume here. We expect that the SC will constrain the presence of the connectivity. However, we do not expect the SC strengths to map directly onto the strength of the precision due to individual differences, as well as variations within individuals across functional brain states. Further, the SC can be expected to have different contributions across frequency bands yielding different connection weights. For these reasons we use the binarized SC to determine the penalization structure. We estimated the penalization values *λ*_1_ and *λ*_2_ using cross-validation as described in the next section (section 2.3.2).

By optimizing the penalized likelihood, we leveraged the information in the SC as a hypothesis for our lasso estimate. The output of the lasso when optimizing Equation 12 is the precision. We derive our graph *G* with vertices *V* = 1, 2, …, 114 and edges *W_est_* = *G_ij_* = 1, *i, j* ∈ *V* from the precision based on the non-zero values in 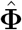. The final precision matrix 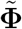 is estimated under the unpenalized Gaussian likelihood for the set of edges ***W**_est_* defined by the graphical model using the function *ggmFitHtf* (PMTK3 toolbox [38]) which optimizes (unpenalized Gaussian log-likelihood):

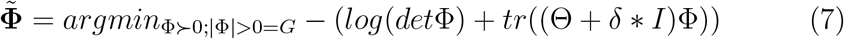

Since **Θ** (covariance) is usually rank deficient, we add a small value (*δ*) along the diagonal to make it full rank. We fixed *δ* as 0.001 times the maximum value along the upper triangle of the covariance.

#### 2.3.2. Cross-Validation

The adaptive graphical lasso provides a way to allow the SC to constrain the statistics of data. However, we wanted to test whether the SC was actually useful as a constraint. To test this, we compared whether the adaptive graphical lasso (which used the SC) produced estimates of the precision that show reduced error relative to applying the graphical lasso. Note that applying the graphical lasso would be equivalent to having the penalization inside and outside the SC be equal i.e. *λ*_1_ = *λ*_2_. We estimated the appropriate value for *λ*_1_ and *λ*_2_ using cross-validation.

We split data into four ensembles, and repeated the following analysis with each ensemble. We estimated the precision 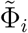 on one ensemble of the data (*i*) and estimated the deviance when using this precision as the inverse for the covariance **Θ**_*j*_ for all the other ensembles *j* of the data (and vice versa). Deviance was estimated as:

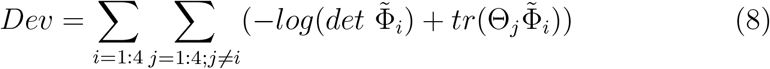

#### 2.3.3. Partial Coherence

In every frequency band, or for each iteration of our simulation, we estimated the precision for complex-valued data incorporating amplitude and phase for a frequency band. The normalization of the precision yields the the partial coherence (*PC*) [19], estimated using:

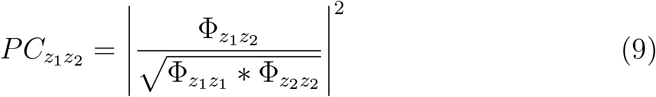

### 2.4. Contemporary Methods for Functional Connectivity

We considered three alternative methods to compare the cGGM model estimated from AGL: coherence, imaginary coherence and the cGGM estimated when regularizing using the L2 norm. Coherence is a commonly used metric for functional connectivity that assesses the consistency of amplitude and phase between two areas in a frequency band. We estimate coherence *C* from the cross spectral density Θ, where *z*_1_, *z*_2_ are two sources, as:

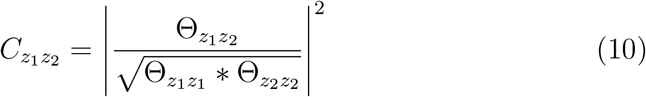

Coherence is a simple summary statistic that captures linear dependence between complex-valued data, however, with M/EEG data, coherence tends to be influenced by volume conduction and/or source leakage. Source leakage is the spread of source activity from one source to neighboring sources due to inexact source localization. Imaginary coherence has been proposed as an easy alternative that is able to reduce the influence of volume conduction and zero-phase lag connectivity (such as would exist from source leakage). The idea is to minimize this effect by estimating the consistency of the imaginary part of the cross-spectral density between two sources. We measure it using (where *imag* refers to the imaginary component of the complex value from the cross spectral density):

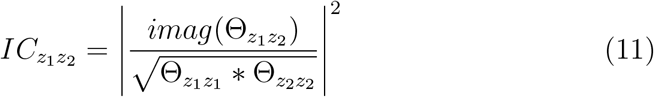

Note that neither of these methods offer an intuitive thresholding heuristic, it must be separately imposed as a posthoc threshold derived from asymptotic distributions [7] or from bootstrapping. In contrast, the AGL allows thresholding of the partial coherence as a part of the process of its estimation. For the coherence and imaginary coherence we derive this threshold from bootstrapping [39] by sampling as many samples as available, with replacement, 1000 times and keeping edges with strengths above the *C* or *IC* value that is at an alpha value of 0.05.

Finally, we consider an alternative regularization to estimate the complex Gaussian Graphical model – an L2 norm penalization. This style of regularization does not push values to zero but instead minimizes them to optimize the likelihood. The penalized likelihood for the L2 norm inverse is:

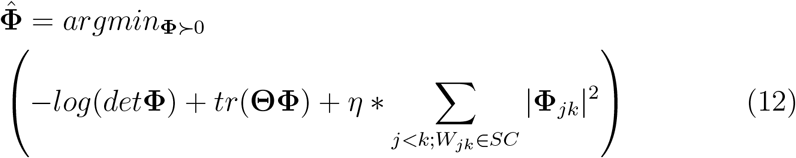

As the L2 norm does not push values to zero, we are forced to identify a threshold for inference on the edges of the cGGM. Using a novel cross-validation procedure that mirrors the approach we applied under the AGL (using the likelihood function to estimate deviance), we optimize for the penalization applied under the L2 (*η*) and the threshold for identifying the edges of the cGGM. The threshold to be applied is determined as a percentile metric of the weights whose optimal value is identified using cross-validation.

### 2.5. Simulations

#### 2.5.1. Overview

The generative model we use in simulations is a complex-valued multivariate normal where the non-zero values in the precision are determined by an undirected network. For each simulation, we generated novel networks with random weights for edges. While the edge locations are consistent within a simulation, we randomized the weights on the edges across simulations. We examined each simulation under two (or more) sampling scenarios while keeping the signal-to-noise ratio constant – one where the number of samples is comparable to the number of nodes and one where there are many more samples than the number of nodes. For each simulation, we examined the ability of the AGL to demonstrate usefulness of the network constraint, to recover the true edges, control the false positives and correctly estimate the edge weights of the cGGM.

#### 2.5.2. SC based simulation

To generate novel precision matrices for each iteration of our simulation, we retained the edge locations from the original SC but simulated random weights for the edges sampled from a normal distribution. Finally, each edge is assigned a random phase (*μ*) based on sampling from a gaussian distribution (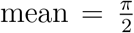, std. dev. = 0.25). After multiplying each edge weight with the phase, we can generate the precision. This represents the complex-valued, circularly symmetric precision matrix (**Φ**) for a frequency band.

Using the precision, we can directly generate the cross spectral density (covariance) as its inverse (**Θ** = **Φ**^*−*1^). As the CSD has a real-valued equivalent [26], treating the real and imaginary components as separate variables governed by the single covariance structure, we sampled complex-valued Gaussian values using the Matlab function *mvnrnd* operating on the equivalent real-valued covariance matrix. The complex-valued samples are used to estimate the cross spectrum (covariance) 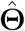.

#### 2.5.3. Fake SC network

We wanted to determine if the AGL could be used to assess the informativeness of a hypothesized network. We begin with the same structure as in the SC simulation above, generating precision matrices and data in the same way. However, we change how we apply the AGL to estimate the cGGM. Rather than provide the true network as a constraint for the AGL, we instead provide a fake network generated by shuffling the nodes of the structural connectome, thus allowing us to preserve the degree distribution of the original network. We shuffle nodes using the *randperm* function in MATLAB. Every iteration of the simulation, we shuffle the nodes of the SC so that the number of edges and general connectome structure are retained while the actual node identities are altered. In this way, we intended to examine whether the AGL permits inference about the hypothesized network, i.e. whether we can we judge how close to the truth the hypothesized network is from the penalizations chosen under cross-validation. We expect that for the fake networks, there is no information in the hypothesized network and thus there should be no difference in penalty inside or outside the fake network.

#### 2.5.4. Forward Solution and Source Localization Simulation

The MEG forward model is an estimate of the magnetic field measured at MEG sensors above the scalp generated by current sources located in the brain. The data analysis presented in this paper makes use of data obtained using the Neuromag MEG system consisting of 306 MEG coils at 102 locations above the scalp (shown in Figure 2.5.4). At each location, there are 3 sensors - one magnetometer that measures the component of the magnetic field passing through the coil and two planar gradiometers that measure the gradient of this magnetic field in two orthogonal directions. For both the simulations and the data analysis presented in this paper, we made use only of the orthogonal pair of planar gradiometer coils (102 pairs of sensors at 102 locations) as planar gradiometer coils have better spatial resolution than magnetometer coils [40].

**Figure 2:**
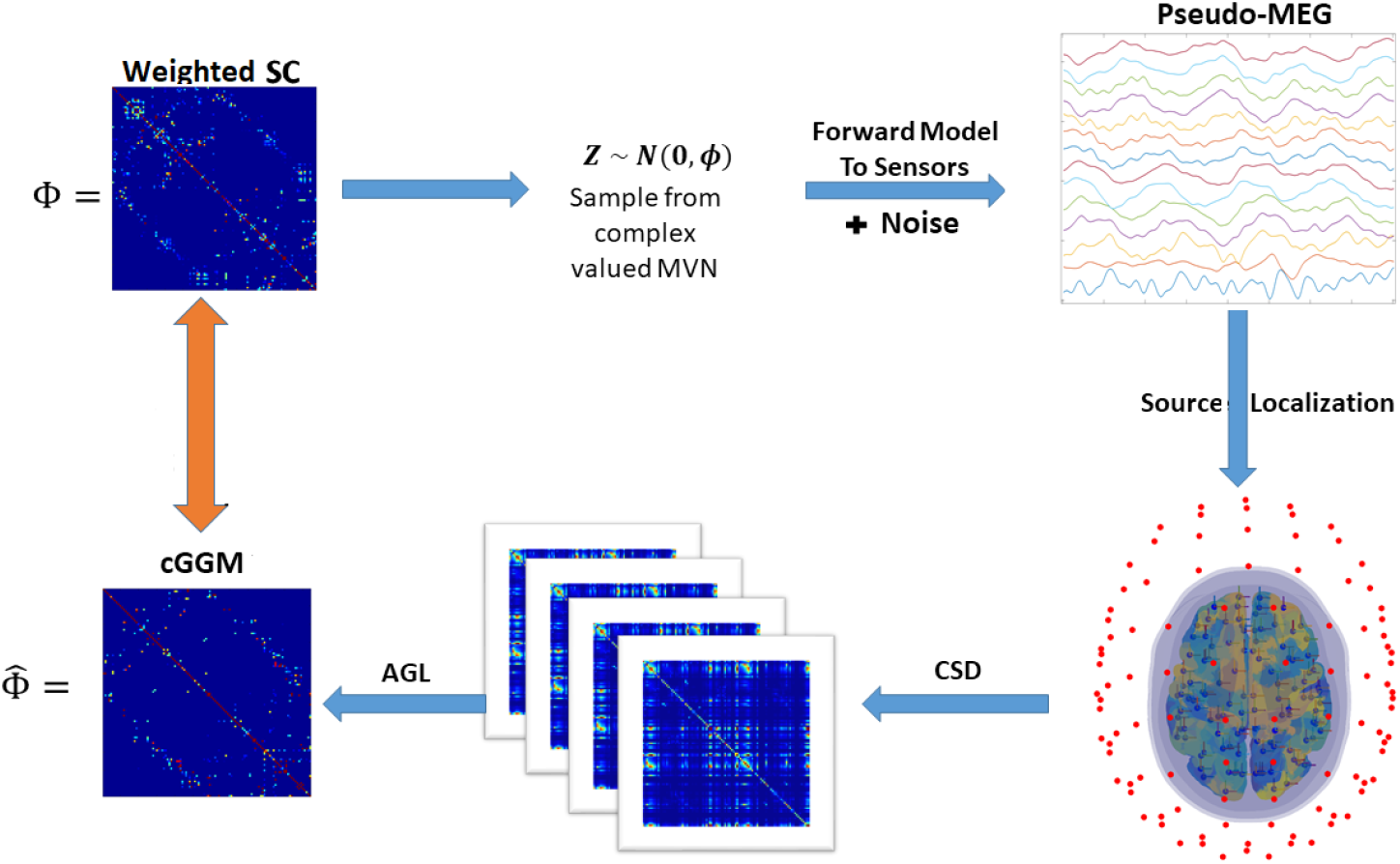
Source Localization Simulation describing each figure above in a clockwise manner from top left: First, we used the SC edge locations to constrain the precision on each iteration of the simulation. We generated random weights and phases for each edge. Second, we sampled from a complex-valued multivariate normal distribution using the precision generated in the first step. Third, we used the MEG forward matrix to forward model the samples to the sensors. Fourth, we applied an inverse solution to source localize data. Fifth, we split the data into 4 ensembles of 120 samples (represented are the 4 covariance matrices from these ensembles of data). Finally, these 4 ensembles served as the input for the adaptive graphical lasso. The estimated precision from this procedure was compared to the original precision (orange arrow) by examining the penalizations applied, the edges recovered and edge weights recovered.

The head model was developed from the *fsaverage* MRI image from the Freesurfer toolbox [41]. The tessellated cortical surfaces for right and left hemisphere were extracted using the *recon-all* pipeline in Freesurfer and then downsampled to 81000 (81k) vertices (*mris decimate* from Freesurfer). We used this surface to constrain dipole orientation and define the volume of the model corresponding to the cortex. We generated the inner skull, outer skull and scalp surfaces approximated with 2562 vertices from the fsaverage head generated using the *mri watershed* function. Using these surfaces, and with the conductivities of the scalp, CSF and brain set at 1 S/m and the skull at 0.025 S/m (i.e., 40 times lower conductivity), we used the OpenMEEG toolbox [42] to compute a Boundary Element Model (BEM) to generate the MEG forward matrix. Each row of the MEG forward matrix is the magnetic field gradient detected across all 204 gradiometers from a unit current density source at one of the 81k cortical surface vertices.

Using the Lausanne parcellation for 114 cortical ROIs [22], we subdivided the cortical surface and identified vertices belonging to each ROI using the volumetric parcellation of the *fsaverage* brain. Using this organization of vertices we then reduced the representation of the current source for each ROI down to a set of 3 dipoles in the *x*, *y*, and *z* directions at a single location. The location of the source for each ROI was selected by taking a weighted average of vertex locations where the weight of each location was determined by the magnitude (L2 norm) of the field generated at the gradiometers. In this way, we reduced our source model to 114 source locations, with 3 sources at each location in the canonical x,y, and z directions. We computed a new MEG forward matrix (***M***) of dimension 204 x 342 using OpenMEEG which approximates the linear mixing of source activity at the gradiometers to generate the measured MEG signals.

We simulate source activity ***S*** across 114 areas using the precision with edges determined by the structural connectome, i.e. one sample from the real-valued equivalent of the inverse of the precision is a 114 x 1 vector. To this source activity, we add independent noise and then forward model it to the MEG sensors using the forward matrix. A sample of the MEG data is represented as a complex-valued vector ***B*** of length equal to the number of MEG sensors (204 sensors). The set of samples of ***B*** relates to source activity ***S*** by:

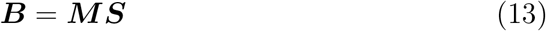

where ***M*** is the MEG forward matrix.

We localize activity to the 342 sources (3 directions - along *x*, *y* and *z* axes at 114 locations) by inverting the reduced lead field using regularized weighted minimum norm estimation (weighted L2 norm [43]) and applying it to data at the scalp. We estimated the inverse ***M**^−^* using (where *ν* is a penalization term):

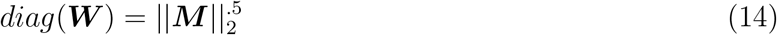

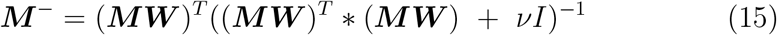

We defined *ν* as the 10th percentile of the the weights of ***MW***. The estimated source activity is then ***S*** = ***M**^−^ **B***. We identify the time series for the three dipoles along the *x*, *y*, and *z* directions. Using a singular value decomposition at each ROI we identified the optimal orientation of the dipole as the first singular vector. Using the first singular vector at each ROI, we reduced the source data from 342 x 1 to 114 x 1 for each sample.

We used the source localized data as the input to the AGL to estimate a cGGM model. We also estimated the coherence, imagina ry coherence and cGGM under the L2 norm. This simulation tests the ability of different methods to overcome the spatial blurring induced by the process of source localization - source leakage and incomplete demixing of source signals. For a visual depiction of this simulation, please see Figure 2.5.4.

#### 2.5.5. Estimating Accuracy of Network Reconstruction

Across all simulations we used the ground truth cGGM to help us understand the performance of different algorithms. To understand the whether the AGL is better than the traditional graphical lasso, we examine the penalization applied on the edges and non-edges of the network provided as a constraint. Across methods, we look at the number of true edges recovered, the number of false positives estimated and the accuracy of connection weights estimated. We calculated the Pearson correlation between the Fisher r-to-z transformation of the ground truth weights with the Fisher r-to-z transformed reconstructed estimate across the set of true edges.

### 2.6. Application to MEG data

#### 2.6.1. MEG Data

The MEG data we analyze was shared by the Cambridge Centre for Ageing and Neuroscience (CamCAN). CamCAN funding was provided by the UK Biotechnology and Biological Sciences Research Council (grant number BB/H008217/1), together with support from the UK Medical Research Council and University of Cambridge, UK. This data was obtained from the Cam-CAN repository (available at http://www.mrc-cbu.cam.ac.uk/datasets/camcan/, [44, 45]).

MEG data was collected using a 306 sensor VectorView MEG system (Electa Neuromag, Helsinki). The 306 sensors consisted of 102 magnetometers and 204 planar gradiometers. The data were sampled at 1000 Hz and highpass filtered at 0.3 Hz. This data was run through temporal signal space separation (tSSS, Taulu et al., 2005; MaxFilter 2.2, Elekta Neuromag Oy, Helsinki, Finland) to remove noise from external sources and to help correct for head movements (location of the head was continuously estimated using Head Position Indicator coils). MaxFilter was also used to remove the 50 Hz line noise and also to automatically detect and reconstruct noisy channels.

#### 2.6.2. Spectral Analysis

We extracted 480 seconds of resting state gradiometer data for a single individual. We first applied a bandpass filter between 0.5 to 100 Hz and a notch filter at 50 Hz to remove line noise. We built elliptic filters (designed using *fdesign.bandpass* function in MATLAB) with stop band set to 0.5 Hz below and above pass band, stopband attenuation set to 100 dB, and pass-band ripple set to 0.02. Bandpass filtering was then done using the *filtfilthd* function in MATLAB to minimize phase distortion. We analyzed five frequency bands: delta (1-3 Hz), theta (4-7 Hz), alpha (8-13 Hz), beta (14-29 Hz) and gamma (30-80 Hz). Within each band we optimized the dipole orientation across 114 ROIs as described in section 2.5.4. Using the bandpass filtered data we were able to estimate adaptively source localized data for each subject and within each frequency band. Source localized broadband data, using band specific source dipole orientations, was multitapered and Fourier transformed in 1 second windows. We used the central frequencies in every band (2, 5.5, 10, 22, and 55 Hz), to avoid averaging over frequencies, to generate a 480×114 complex-valued matrix used for estimating the cross-spectral density. Additionally, we compared results when using other frequencies within a band as well to ensure no large variability existed at any individual frequency within a band relative to another.

Using the complex valued data withing each frequency band, we have a 480×114 matrix which served as the input for estimating the cGGM. We split the 480 samples from 114 sources into 4 continuous ensembles of 120 samples each based on the expectation that we would have robust, stationary networks estimable with 120 seconds [46]. Further, having 4 ensembles allowed for 4-fold cross-validation. Within each ensemble we estimated the cross-spectral density and using the adaptive graphical lasso, the precision. We then followed the same procedure as described earlier in the section 2.3.2 on cross-validation. Thus, we had at the end of the analysis for each subject, cGGMs across all 5 frequency bands.

### 2.7. Code Availability

We make all code and data used in the analyses in this paper available at https://github.com/wodeyara/cGGMs-for-Coherence.

## 3. Results

We present several simulations to demonstrate the ability of Adaptive Graphical Lasso (AGL) to estimate a complex Gaussian Graphical Model (cGGM). We first demonstrate a proof of concept by recovering a small network generated from a cGGM. We then scale up the model to a cGGM whose graph is defined by the structural connectome obtained from a probabilistic DTI atlas [21]. When we simulate from this model, we show that the AGL can recover the graph with high accuracy as the number of samples is increases. We then demonstrate that this method can infer when the hypothesized network used for the AGL contains information relevant to the generation of the observed data. We compare the AGL estimation of a cGGM with contemporary methods for functional connectivity and show how the cGGM estimation outperforms these contemporary approaches at network recovery. Using a forward model and source localization, we show that cGGM estimation outperforms contemporary algorithms at network recovery following source reconstruction. Finally, we estimate cGGMs of source localized MEG data and find that different frequency bands show different resting state connectivity profiles within the structural connectome.

### 3.1. Simple Five Node Network

As a proof-of-concept simulation, we examined network recovery of a sparse five node network with five connections between nodes. The network is defined by a complex-Gaussian Graphical model (cGGM) represented by a complex-valued matrix - the precision (inverse of the cross-spectral density matrix at a frequency/in a frequency band). When there is no connection (edge) between a pair of nodes the entry for that pair in the precision matrix is zero and, when a pair is connected, the precision is a complex-value defining the strength and relative phase difference between two nodes (see Figure 3A). The precision expresses the linear dependence of the complex-valued (amplitude and phase) data at the two nodes, after removing the (linear) influence of all other nodes. In a cGGM, this precision matrix contains all the parameters of our generative model. We sample the observed data for each node from the inverse of the precision, the cross-spectral density, under a zero-mean multivariate normal distribution. We apply the Adaptive Graphical Lasso to the observed data using knowledge of the network structure but no edge weight information and placing no constraints on the penalization applied. We consider two case, one where we have a small number of samples (24 independent samples) and second, when we have a large number of samples (240 independent samples). Each simulation (24 and 240 samples) is repeated 200 times.

**Figure 3:**
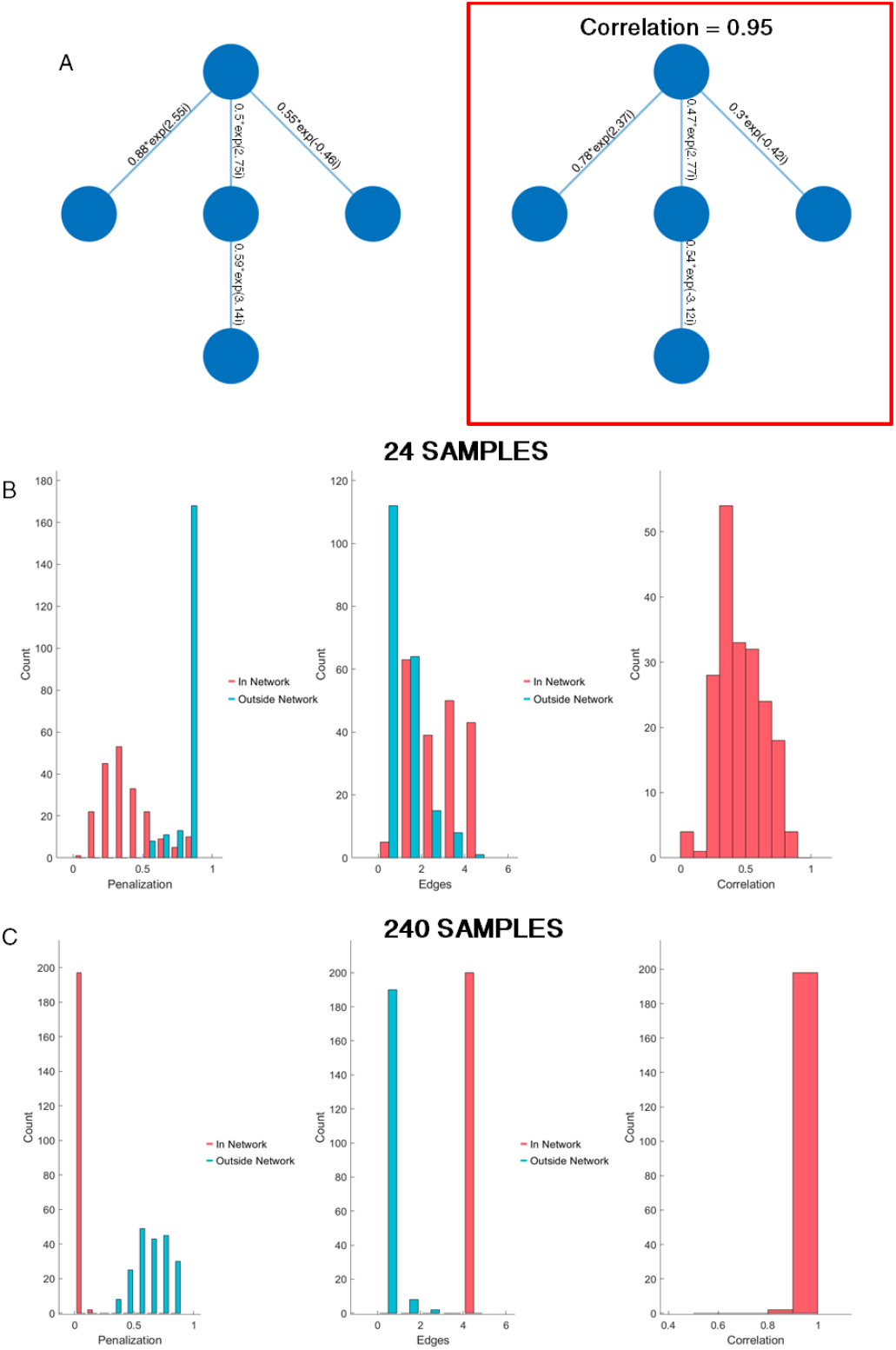
AGL provides accurate Five Node cGGM inference. **(A)** (Left) We simulate from a simple 5 node network with complex valued edges. (Right) An example recovered network with weights nearly perfectly reconstructed (using 240 samples). **(B,C)** We test the ability of the AGL to recover the correct cGGM through three metrics. The first (left column) is a bar plot of the penalization applied inside (red) and outside (blue) the network provided as a potential constraint. This penalization is chosen based on cross-validation. The penalization outside the network is consistently higher than inside the network. In the middle we show how well correct edges are recovered (red) and the number of incorrect edges (false positives) are generated. On the right we show the correlation between the correct edges in the recovered model with the true edge weights. The adaptive graphical lasso is able to identify that the network provided as a potential constraint is useful (penalization applied in network < penalization applied outside network), controls the false positives and recovers network weights reasonably well, even at 24 samples (B). When 240 samples (C) are available, we find that the AGL recovers the network almost perfectly. In the middle column we see that all 5 edges are recovered and the edge weights are nearly perfectly reconstructed with the correlation near 1.

We examine the usefulness of several metrics to assess the accuracy of the estimate of the network graph using the AGL. First, we expect that the penalization parameters from the AGL will demonstrate the usefulness of prior knowledge of the network. The AGL should place lower penalization on the edges in the network and higher penalization on the edges outside the network. We see from the penalization distribution (Figure 3) that there is reduced penalization for true edges relative to non-edges, as we expected, both with a low and high number of samples. As the search process allows the AGL to place the same penalization everywhere, the penalization values assess the usefulness of our prior knowledge of the graph. Our second metric of interest is edge recovery - how many true edges and how many false positives are discovered? In Figure 4B we can see that the false positives are well controlled (with the distribution concentrated at 0 edges) while we recover between 2 to all 5 of the true edges present when only 24 samples are available. With 240 samples (Fig 3C), we recovered all true edges in the all 200 simulations and nearly always avoided any false positives (95% of simulations). Our final metric of interest is the recovery of the actual weights of the edges - the complex values representing connection strength and relative phase. We estimate this correspondence using a correlation between the true edges and the recovered edges. A high correlation implies that the complex valued vectors tend to align with the orientations and strengths of the original complex valued vectors and a correlation close to 0 indicates incorrect weight and orientation (an orthogonal vector or a zero vector). From Figures 4B and 3C we can see that correlation is around 0.5 with 24 samples while it is nearly 1 with 240 samples. We conclude that we are able to recover weights and edges of the cGGM even when we have only 24 samples, but with (an order of magnitude) more samples, we are almost able to recover the cGGM perfectly.

**Figure 4:**
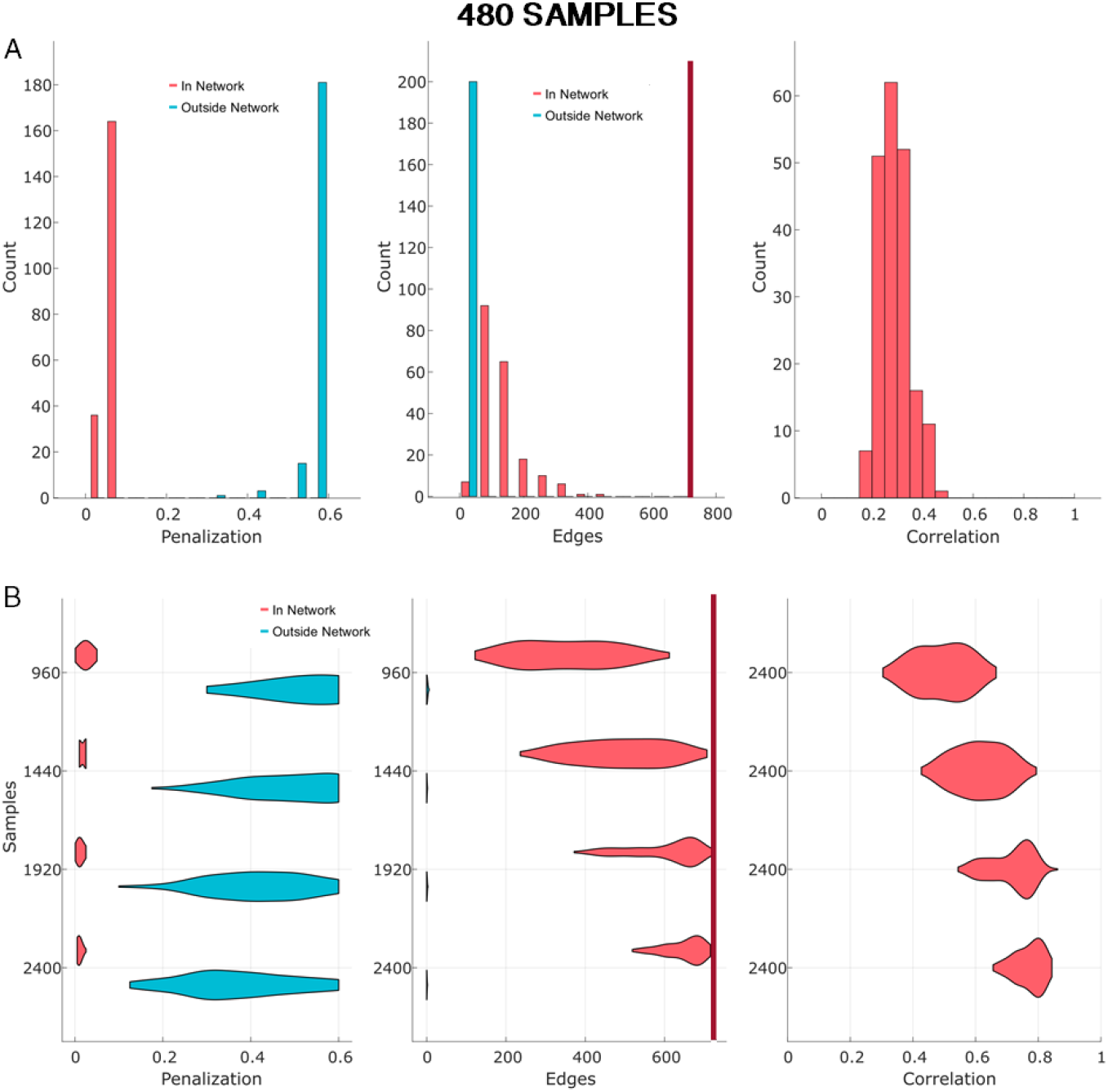
AGL recovers the Structural Connectome cGGM. **(A)** When simulating data from the structural connectome, even in a low samples environment (480 samples), AGL identifies the correct penalty structure (left column) and controls false positives (middle). However, recovery of edge weights (right) is poor. **(B)** We show changes in performance as a function of the sample size. As the number of samples increases, AGL improves significantly, with network recovery almost perfect at 2400 samples. In middle column red vertical line represents the total number of true edges present.

Note that as we have access to ground truth in these simulations we are able to understand performance through edges recovered and correlation with original edge weights. But even without this information in real data, we can still assess the whether prior knowledge of the network meaningfully contributes to the cGGM by examining the structure of the penalty.

### 3.2. Recovering the Structural Connectome

In the second simulation, we considered an order of magnitude increase in the number of nodes and edges. To define the true edges of the network, we use a structural connectome estimated from HCP-842 data, organized by the Lausanne parcellation of 114 cortical regions. This network is sparse, with 720 edges out of a total possible of 6441 edges. By simulating random weights and phases for the complex values at each SC edge, we generate independent realizations of the precision for each iteration of the simulation. The inverse of the precision could represent the cross-spectral density estimated from intracranial electrocorticography (ECoG) sampling the entire cortex. Similar to the first simulation, we examine the performance of the the AGL to estimate the correct cGGM when we have 480, 960, 1440, 1920 and 2400 samples at each of 114 nodes. Under each sampling scenario, we simulated 200 realizations of a cGGM with different precision values at the edges corresponding to the connectome.

Network recovery under AGL in a high sampling situation (2400 samples) is nearly perfect (Figure 4). The penalization structure indicated lower penalization on SC edges relative to non-SC edges, the false positives are controlled and real edges identified (≤500 of 720) and finally, the edge weights are well recovered (correlation ≥ 0.7). This showed that AGL is able to establish a penalization structure that demonstrates the guiding presence of the structural connectome. Even when we simulated only 480 samples the AGL minimized false positives, showed the usefulness of knowledge of the SC and reasonably recovered the network weights. We conclude that low numbers of samples do not pose an impossible hurdle in recovering the structural connectome under the AGL and further, that large, sparse networks are recoverable with the AGL.

### 3.3. Network Recovery under a Fake Network

In empirical data we usually do not have access to the true underlying network. We can use anatomical knowledge drawn from reasonable estimates (for e.g. DTI as in the HCP-842 dataset). However, when applied to an individuals M/EEG data we cannot be sure of how much error is present. We test if we can infer, from the AGL, whether a given network is informative of the true underlying generative network i.e. to force model mis-specification and test its effects. We expect that the penalization structure of the AGL will reflect when we have used an incorrect network as a potential constraint. An alternative hypothesis is that the AGL consistently under-penalizes the network provided to it, regardless of the true generative network, thus making the prior network critical and making inferences on whether the hypothesized network is appropriate impossible. We test these hypotheses by shuffling node identities for the network used by AGL on every iteration of the simulation. However, we generate data from the true, unshuffled precision with non-zero edges at the structural connectome edges.

Examining the penalization structure (Fig 5B), we find that the AGL does not place lower penalization values on the fake network edges, instead having a similar penalization across the network (as we expected). However, this does not imply network recovery in either the fake or the true networks (Fig 5B and 5C, right column), with both the false positives and the true edges suppressed in both networks at 480 samples. However, at 2400 samples, we see that while the penalization again is similar inside and outside the fake network, more true edges are now estimated (Figure 5C, right), while the false positives driven by the fake network continue to be controlled (Figure 5C, middle). This suggests that while the AGL remained constrained by the network provided, (1) an incorrect network can be inferred from the penalization structure and (2) with sufficient samples the true network can be partially recovered.

**Figure 5:**
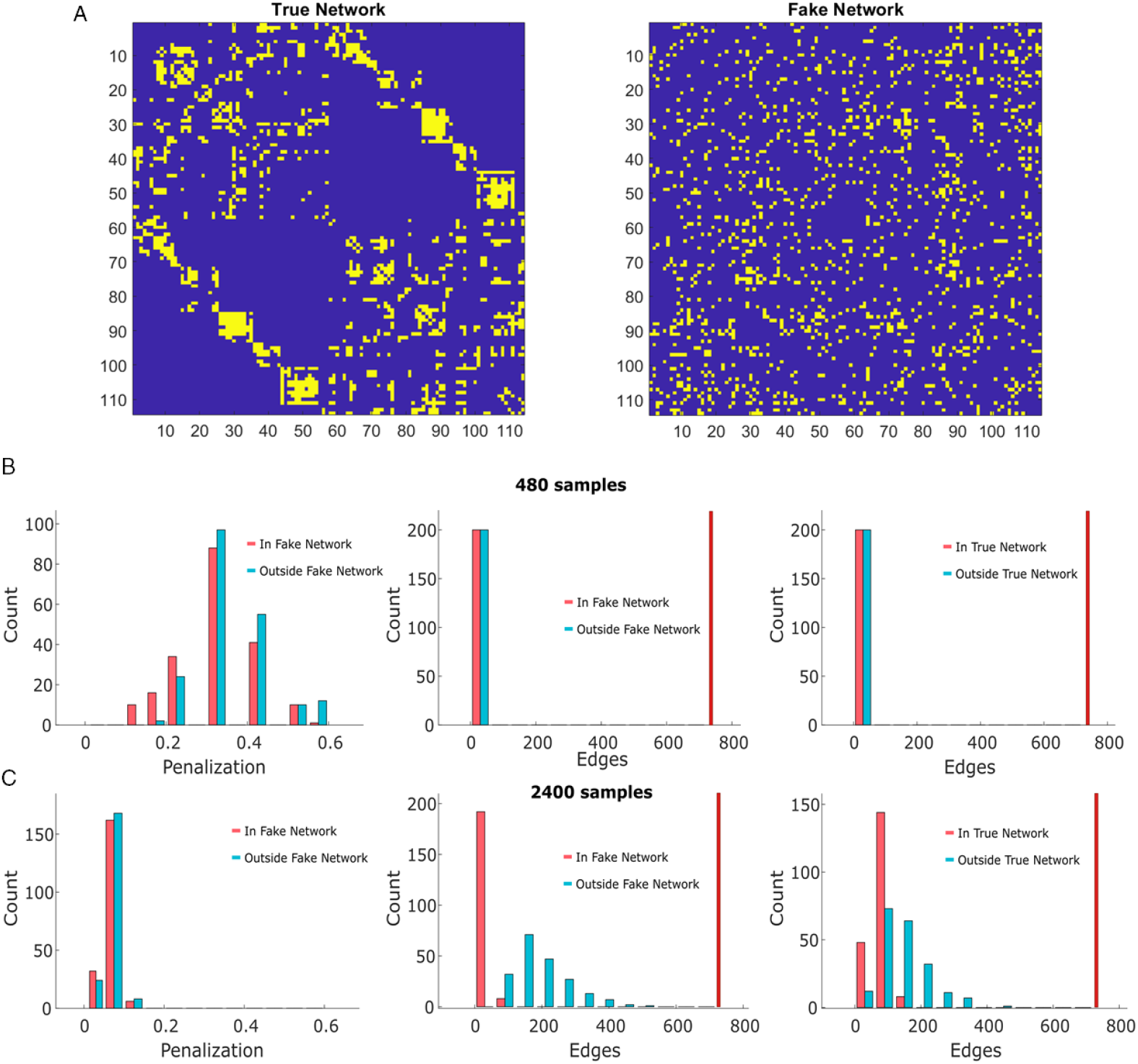
Fake Network provides no useful information. **(A)** The true network represents the cGGM from which the data is simulated. In the fake network, we shuffle nodes in the true network, thus retaining the degree distribution of the original network. We give this fake network as the hypothesis for the AGL. **(B)** In the low samples scenario, we find that the AGL is unable to use the fake network and shows this in the penalty structure (left column) with the penalization inside and outside the fake network set similarly high. Due to the high penalization it is unable to identify any SC edges (right column) but also controls false positives from the fake network (middle column). **(C)** At higher numbers of samples, the AGL is able to partially overcome the fake network. Despite setting penalization inside and outside the fake network at around the same value (left column), a considerable number of true edges are recovered (left column) with the caveat that several false positives also result. Note that there can be overlap in the edges in the fake network with edges in the true network. As before, the vertical red line represents the total number of edges present in the model.

### 3.4. Comparing AGL to Contemporary Network Recovery Approaches

Many contemporary algorithms aim to estimate the network underlying EEG/MEG/ECoG data. Several studies have attempted to use coherence/imaginary coherence to estimate the influence of the structural connectome [47, 12] on the networks recovered. We compared three methods that make similar assumptions about the data as a cGGM - coherence, imaginary coherence and the partial coherence estimated under an L2 norm inverse [48].

We first compared these methods when recovering a network with structural connectome edges when estimated from 480 samples. We implemented an inference technique for coherence and imaginary coherence that relies on bootstrapping (see *Methods, section 2.4*), and we use a cross-validated deviance metric to infer the correct L2 norm penalty to apply when estimating the partial coherence. We compared the methods on the true positives, false positives and the network weight recovery (Figure 6).

**Figure 6:**
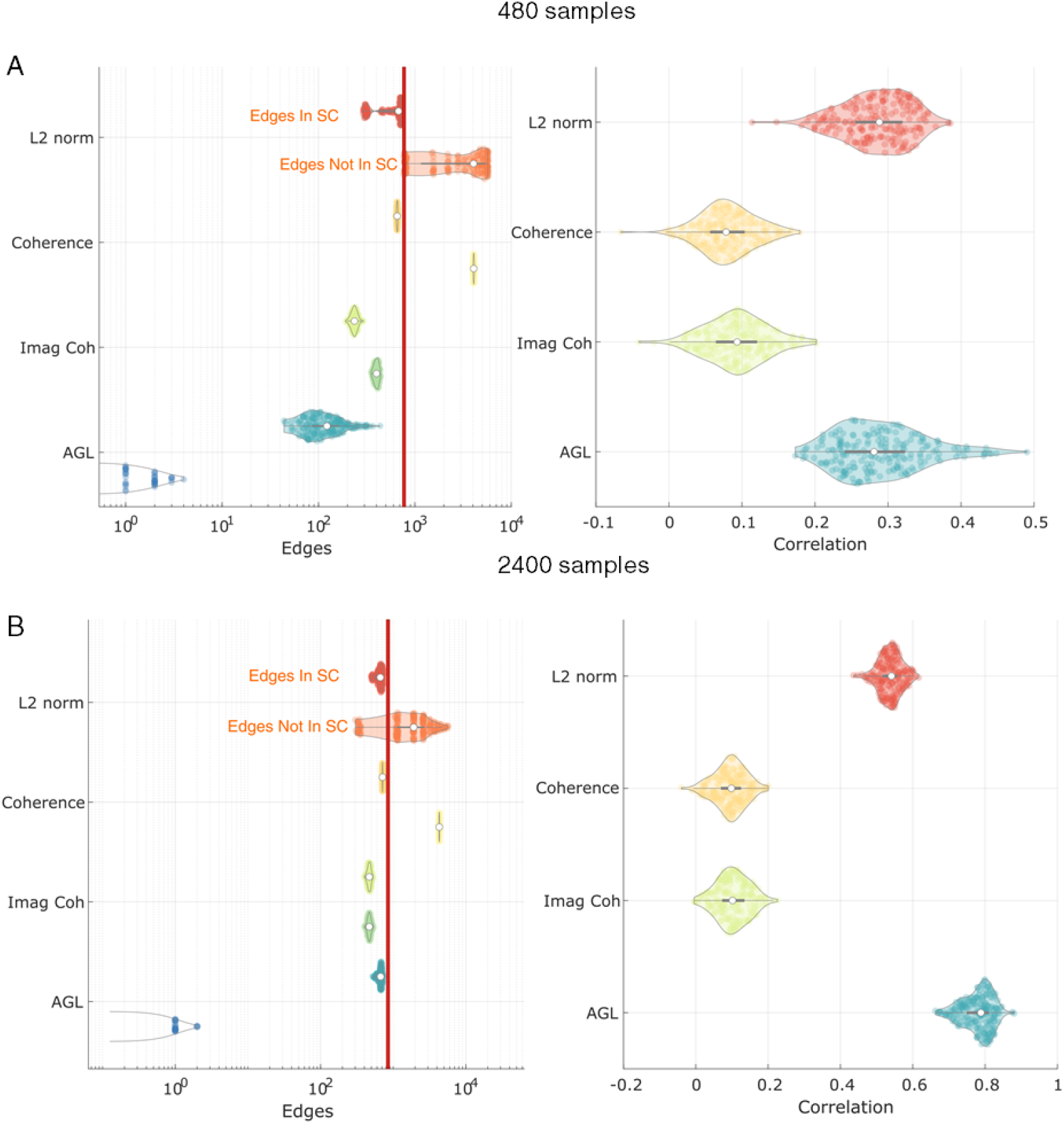
AGL outperforms alternative methods at cGGM recovery. We compare AGL to an L2 norm inverse, coherence and imaginary coherence. We show two cases: **(A)** 480 and **(B)** 2400 samples. **(A)** (Left) For each method on the y-axis, there are two violin plots with the upper violin representing edges in the SC and the lower violin showing edges outside the SC. No method controls the edges outside the SC (false positives) as well as the AGL (see bottom violin), however, all alternative methods recover more edges in the SC. The L2 norm inverse recovers edge weights comparably to the AGL. **(B)** At 2400 samples, the adaptive graphical lasso outperforms all other algorithms at controlling both false positives and at edge weight recovery. Note that since the L2 norm is thresholded based on percentiles of edge weights, the number of edges recovered outside the SC can be discrete.

**Figure 7:**
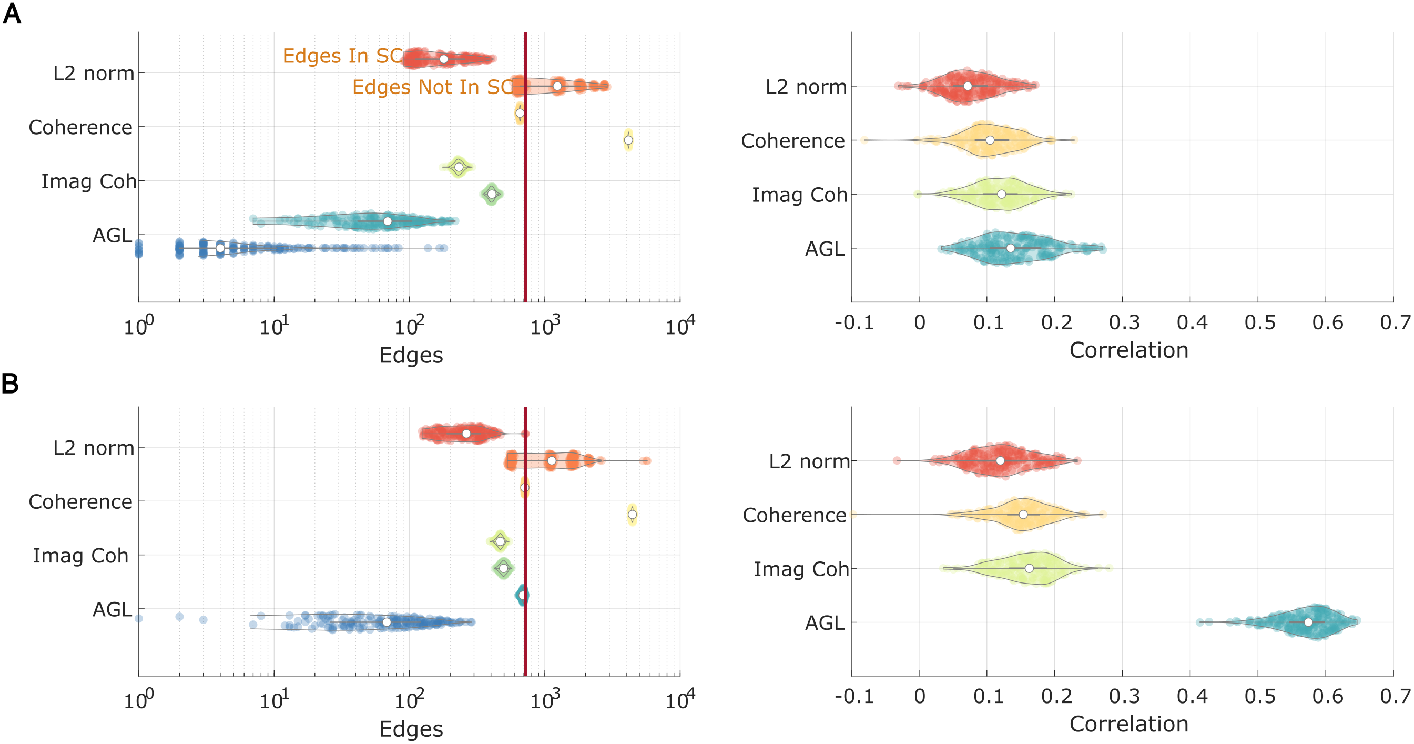
AGL outperforms alternative methods at cGGM recovery under Source Localization. We generate data from a cGGM in the brain with SC edges and forward model to MEG sensors. After applying source localization (weighted L2 norm inverse), we attempt to recover the original network. **(A)** At 480 samples, while AGL controls false positives better than other algorithms (right), all methods are comparable at cGGM edge weight recovery. **(B)** AGL outperforms all algorithms at 2400 samples at recovering true edges, controlling false positives and recovering edge weights.

**Figure 8:**
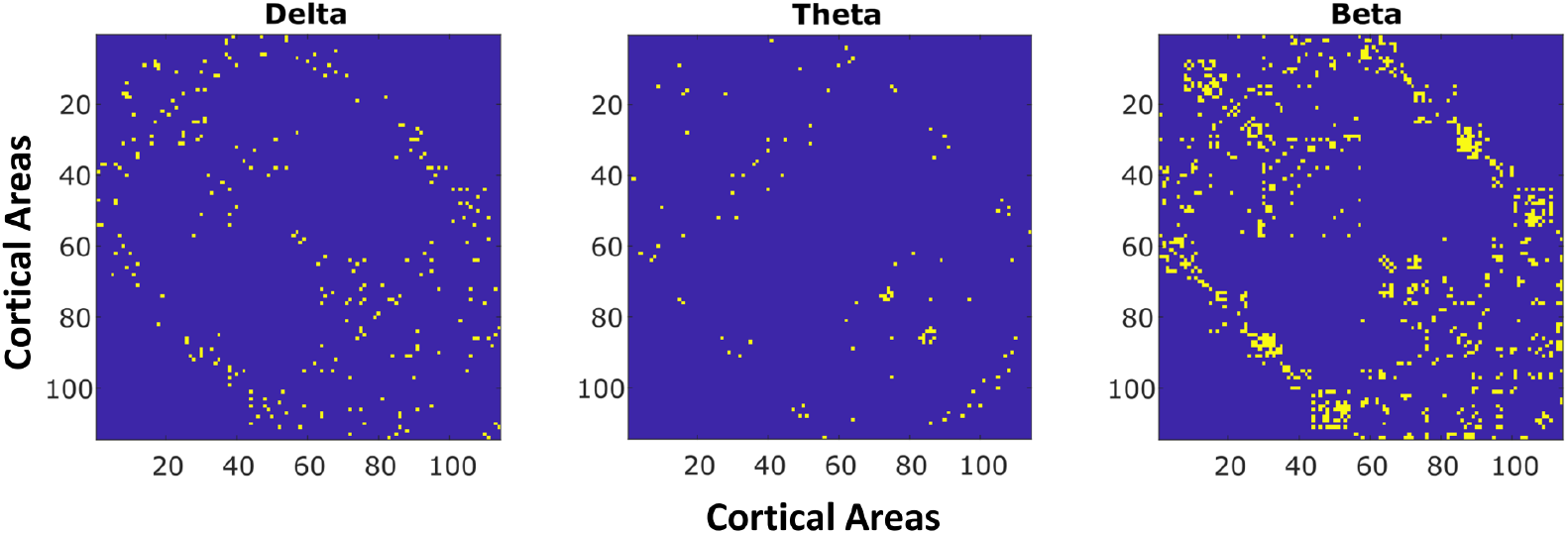
AGL recovers Different SC constrained cGGMs in different frequency bands. We apply the AGL to delta, alpha, theta, beta and gamma frequency bands. We show here results from the bands where there was a lower penalization applied to the SC. In the Delta, Theta and Beta bands we show the edges that were maximum in strength for that band. This selection shows how there exist different networks at each frequency with the beta band showing the greatest spread across the SC and the theta band network showing the greatest selectivity.

We found that, at 480 samples, all methods were able to recover the true SC edges, however, they also estimated a considerable number of false positive edges. The AGL was considerably better at controlling false positive edges than all other methods, with the imaginary coherence performing better than coherence and the L2 norm. When estimating network weights however, the L2 norm inverse and the AGL did considerably better than the coherence and imaginary coherence. When examining 2400 samples, we saw similar performance differences across methods, with the AGL continuing to outperform all other methods at controlling false positives. Further, the AGL is better than the L2 norm inverse at estimating the network weights of the true network at 2400 samples. We conclude that the AGL outperforms contemporary algorithms at recovering the underlying network, both when we have limited samples and when we have large numbers of independent samples.

### 3.5. Network Recovery under Source Localization

In M/EEG research, we must both recover the network from limited samples and reduce the impact of signal leakage from source localization. We simulate sources similarly to earlier cases: by sampling from the inverse of the precision which has complex values at the non-zero edges of the SC. This activity at each node is forward modeled using a head model (see *Methods* Section 2.5.4) and get mixed together at the MEG gradiometers. We apply a commonly used source localization technique (weighted L2 norm inverse, see details in *Methods*, section 2.5.4) to attempt to recover sources. We apply the AGL and other algorithms to this recovered source activity.

Examining the 480 sample case, we saw that the AGL continued to out-perform all other models at controlling false positives in the network. However, other network recovery techniques were comparable in recovering true SC edges, with the coherence recovering all edges but also included a large number of false positives. All methods were comparably poor at recovering the network weights, likely due to the inability of source localization to fully separate independent sources (source leakage). When we have more samples (2400), we see that the AGL clearly outperforms all algorithms in all metrics measured, with the correlation with network weights reaching 0.58. This suggests that when more samples were available we were able to partially overcome the problem of source leakage using AGL.

### 3.6. Application to MEG Data

We extracted 480 seconds of preprocessed resting state MEG data from a single subject from the open source CAMCAN dataset. We source localized this data (using weighted L2 norm [43]) to the 114 areas of the Lausanne parcellation. After source localization, we used 1 second windows to get amplitude and phase samples at each frequency from 1 to 50*Hz* using the multitaper method. We applied the AGL with our cross-validation procedure (see section 2.3.2) to estimate the cGGM. Note that since we examined only a single subject, we intended this to only be a demonstration of how the cGGM could be used. Further, we don’t have a ground truth in this situation so we focus on the penalization structure to infer if the structural connectome (SC) is useful information in modeling the coherence. We did find that the SC did serve as a useful constraint (defined as the penalization outside the SC being higher than inside the SC) in the delta (2/3 frequencies), theta (1/4 frequencies) and beta bands (11/15 frequencies), but not in the alpha (0/6) or gamma (0/20) bands. For the cases when the SC was a useful constraint, we examined the edge weights in the delta, theta and beta bands, looking for which band had the highest weight at each SC edge. We show this in Figure 3.6. We can see that the beta band tended to have connections distributed across the cortex, while theta and delta connections are more circumscribed. Delta band showed connections within frontal and cingulate regions and from frontal/cingulate to parietal regions. Theta band showed consistent connectivity across left and right hemispheres between temporal and parietal/occipital regions. Beta band connectivity dominated throughout the rest of the structural connectivity, with little specificity. The absence of evidence for the structural connectome in the alpha band, may reflect our restriction of the structural connectome to cortico-cortical connections as deeper sources (e.g., thalamus) cannot be easily estimated from spontaneous MEG. These thalamocortical connections may be misattributed in our model, an effect expected to be strongest in the alpha band [49]. An-other possibility is that the mechanism of alpha rhythm generation is wave propagation [8]. We conclude that the AGL can be applied to empirical data to discover networks in different frequency bands.

## 4. Discussion

We developed a complex-valued Gaussian graphical model (cGGM) of MEG coherence that can be constrained by knowledge of anatomical connectivity in the structural connectome. We showed that we can infer the parameters of the cGGM using the Adaptive Graphical Lasso, and that this approach performs better than contemporary methods of functional connectivity estimation when we want to take advantage of our knowledge of the structural connectome. In resting state MEG data we showed that estimated cGGMs show differences in connectivity across different frequency bands.

### 4.1. Models of M/EEG/ECoG Functional Connectivity Generation

Considerable work has been dedicated to proposing models of MEG functional connectivity. This works seems to occur along a spectrum ranging from complete biophysical accuracy to purely statistical models. On the biophysical end, mean field modelling techniques are used to understand the functional connectivity [50, 51]. On the statistics end, there is work on fitting diffusion models to the functional connectivity [47]. In between are models that look at the harmonics of the anatomical connectivity [52]. cGGMs stand on the statistics end of this spectrum and serve as a very simple model of how M/EEG or ECoG functional connectivity can arise: as a stochastic process correlated by the structural connectivity. As such, this modeling framework offers a simple hypothesis for data generation that can be compared to current alternatives.

### 4.2. Comparison to Alternative Methods

The main improvement visible from the development of the AGL estimation of cGGMs is the orders of magnitude improvement in the control of false positives. Coherence is particularly affected by false positives, likely from leakage effects and from polysynaptic connectivity. A common method to reduce the influence of leakage effects that occurs when estimating coherence is to use the imaginary coherence [53] or (similarly) to remove all zero-lag connectivity [54]. These approaches operates on the (valid) assumption that there is field spread only at zero lag under pseudo-static assumptions. However, three notable problems with these approaches are:

1. They may be too conservative an estimate and choose to lose considerable information about genuine connectivity in the data.
2. They are still susceptible to spurious connections when genuine long range connections exist at a delay (and activity at these areas is leaked to neighboring regions) [55].
3. Finally, common input driven connectivity persisting through a multi-step path over the SC occurring at a delay is unaccounted for.

In contrast, cGGMs do not ignore zero-lag connectivity and by removing common information that exists at a delay (by operating in the complex domain and using phase information) it avoids the second and third issues above. In our results, we showed the consequences of these issues and demonstrated how the AGL estimate of the cGGM is better than using the imaginary coherence and coherence. Further, this makes a cGGM an optimal starting point for graph theoretical analyses as the definition of a path is made significantly clearer when accounting for the issues listed above.

### 4.3. Using Larger Numbers of Samples in Functional Connectivity Research

All simulations we performed suggested that the number of samples used is a strong determinant of the accuracy of cGGMs of MEG coherence. While we were able to recover a subset of the network when the number of samples remained comparable to the number of edges and nodes, it was clear that there was considerably improved performance with higher numbers of samples. While past work has suggested that for coherence, there can be convergence with a few hundreds of samples [46], we saw that for the imaginary coherence and for cGGMs, larger numbers of samples significantly improved performance. This knowledge provides impetus to use larger recordings to estimate resting state electrophysiological functional connectivity, similar to recent work in fMRI research [56].

### 4.4. Limitations

In the simulations we assumed a generative model where different parts of the brain show random oscillatory behavior linked by the structural connectome. This could be represented using a zero mean complex multivariate normal with a circularly symmetric precision. More detailed mean field models of neural activity may be more phenomenologically accurate [50], although, past work suggests there is limited gain in using them when explaining empirical data [47, 57]. As such, there is value in having multiple models to explain the data as a function of the question being posed.

While cGGMs offer a clearer definition of a direct connection between areas, this does not imply that these are the only important connections. As suggested by the notion that coherence supports communication in the brain, coherent connections could have important shared information that is considered redundant under the cGGM framework. As such, the choice of using cGGMs to analyse data could be a function of the questions being posed or they may be better used in concert with coherence as proposed by [15].

In humans, structural connectivity is only estimated from Diffusion Weighted Imaging (DWI) and is an imperfect measure, subject to its own limitations [58, 59]. There are difficulties in tractography linked to overlapping fiber bundles that make it hard to identify correct bundle endpoints and strict correction of incorrect streamlines can rapidly lead to large numbers of false negatives [59]. Our decision to remove non-homologous inter-hemispheric connectivity may also have introduced a few false negatives. Finally, we used a group-averaged SC template for all the subjects, and while individual variability in SC is low [60], better models may be built using an individualized SC estimate.

Source localization can be formulated in several ways based on prior assumptions. While we used a weighted *L*^2^ norm inverse, beamformer reconstruction approaches are also quite common in MEG [61, 62] and require investigation within this framework. Bayesian techniques accounting for priors more explicitly can afford better source reconstruction [63, 64]. Examining these alternative approaches was beyond our scope. Additionally, we chose to limit our analysis to a case where there are 114 sources, a future extension to this work might examine cases with more (fewer) sources. We also ignore for our purposes subcortical source activity and connectivity. This may have led to the large variation in the estimated results in the MEG data. Estimation of subcortical activity in MEG while possible is difficult without explicit prior knowledge [65], and would also potentially benefit from including the magnetometer recordings and developing individual subject head models.

### 4.5. Conclusion

Understanding the relevance of different potential constraints (source modeling/structural connectome) on a “big data” measurement technique such as MEG data improves our ability to infer genuine signal variability from noise. Our work developed a simple model derived from the constraint of the structural connectome and demonstrated that we can recover the parameters for this model in simulations. Our method is useful in clinical situations and for cognitive neuroscience in doing hypothesis driven research. For example, estimating a gamma band cGGM in a working memory task to understand which structural connections are most strongly activated. Another example, as we have recently demonstrated in fMRI research [32], is to examine the influence of lesions and concomitant structural disconnection on MEG functional connectivity. Interpreting M/EEG coherence is contingent on building and comparing different models of the data and we believe our work takes us a significant step in this direction.

